# Antibiotics affect the pharmacokinetics of n-butylphthalide *in vivo* by altering the intestinal microbiota

**DOI:** 10.1101/2024.01.12.575425

**Authors:** Xiangchen Li, Xiaoli Guo, Yixin Liu, Fefei Ren, Shan Li, Xiuling Yang, Jian Liu, Zhiqing Zhang

## Abstract

**Objective:** N-butylphthalide (NBP) is a monomeric compound extracted from natural plant celery seeds, whether intestinal microbiota alteration can modify its pharmacokinetics is still unclear. The purpose of this study is to investigate the effect of intestinal microbiota alteration on the pharmacokinetics of NBP and its related mechanisms.

**Methods:** After treatment with antibiotics and probiotics, plasma NBP concentrations in SD rats were determined by high-performance liquid chromatography-tandem mass spectrometry (HPLC-MS/MS). The effect of intestinal microbiota changes on NBP pharmacokinetics was compared. Intestinal microbiota changes after NBP treatment were analyzed by 16S rRNA sequencing. Expressions of CYP3A1 mRNA and protein in the liver and small intestine tissues under different intestinal flora conditions were determined by qRT-PCR and Western Blot. KEGG analysis was used to analyze the effect of intestinal microbiota changes on metabolic pathways.

**Results:** Compared to the control group, the values of C_max_, AUC_0-8_, AUC_0-∞_, t_1/2_ in the antibiotic group increased by 56.1% (*P<*0.001), 56.4% (*P<*0.001), 53.2% (*P*<0.001), and 24.4% (*P*<0.05), respectively. In contrast, the CL and T_max_ values decreased by 57.1% (*P<*0.001) and 28.6% (*P*<0.05), respectively. Treatment with antibiotics could reduce the richness and diversity of the intestinal microbiota. CYP3A1 mRNA and protein expressions in the small intestine of the antibiotic group were 61.2% and 66.1% of those of the control group, respectively. CYP3A1 mRNA and protein expressions in the liver were 44.6% and 63.9% of those in the control group, respectively. There was no significant change in the probiotic group. KEGG analysis showed that multiple metabolic pathways were significantly down-regulated in the antibiotic group. Among them, the pathways of drug metabolism, bile acid biosynthesis and decomposition, and fatty acid synthesis and decomposition were related to NBP biological metabolism.

**Conclusion:** Antibiotic treatment could affect the intestinal microbiota, decrease CYP3A1 mRNA and protein expressions and increase NBP exposure *in vivo* by inhibiting pathways related to NBP metabolism.

## Introduction

The intestinal microbiota is a complex system. It is an essential part of the intestinal mucosal barrier and plays a vital role in maintaining the dynamic balance of microecology in the organism [1]. Unbalanced intestinal microbiota or disruption of the dynamic balance of intestinal microbiota may bring various problems to the body at the macro level and contribute to multiple diseases [2, 3]. On the micro level, it can also directly or indirectly change the efficacy and toxicity of drugs by affecting the expression of cytochrome P450 enzymes (CYP450) in the host body [4]. Oral antibiotics can induce intestinal microbiota depletion and down-regulate the expression of the CYP450 enzyme, while probiotics can restore enzyme activity and alter substrate metabolism [5]. The complex relations between the host, bacteria, and drugs affect the intestinal microbiota, leading to poor drug efficacy, adverse reactions, and drug-drug interactions in clinical practice [6–8].

N-butylphthalide (NBP), also called butylphthalide, is a monomeric compound extracted from natural plant celery seeds. It is rapidly absorbed after oral administration. NBP is mainly metabolized by CYP3A4 in the liver. Metabolites are conjugated with glucuronic acid and excreted in urine [9]. NBP can inhibit brain tissue damage and promote the absorption of inflammatory factors, and it is recommended to treat acute ischemic stroke as a neuroprotective agent [10, 11]. Changes in autonomic nervous activity and mucin products caused by brain injury can induce specific changes in the intestinal microbiota, and ischemic stroke can induce intestinal microbiota imbalance [12]. Whether intestinal microbiota alteration can modify the pharmacokinetics of NBP is still unclear, and further in-depth studies are warranted.

In this study, we used antibiotics and probiotics to change the intestinal microbiota composition. The effects of intestinal microbiota alteration on CYP3A1 expression in the small intestine and liver and the pharmacokinetics of NBP in SD rats were investigated using 16S rRNA and KEGG analyses. The results lay the foundation to clarify the effect of the intestinal microbiota on drug metabolism and related mechanisms.

## 1 Materials

### 1.1 Instruments

The following instruments were used: high-performance liquid chromatography (HPLC) (Shimadzu, LC-20AD), SCIEX mass spectrometer (MS) (AB SCIEX, API 4000+), PCR amplification instrument (ABI, 2720), enzyme labeling instrument (BioTek, FLX800T, electrophoresis apparatus (Beijing Liuyi Instrument Factory, DYY-6C), a gel imaging system (Beijing Bijing Biotechnology Co., LTD., BG-gdsAUTO130), Nanodrop UV quantitative system (Thermo Fisher Scientific, NC2000), Sequencer (Illumina, Novaseq6000), and fluorescence quantitative PCR instrument (Bio-rad Corporation, CFX).

### 1.2 Drugs and reagents

The following were used: vancomycin hydrochloride for injection (VIANEX S.A.), bifidobacterium quadruplex viable tablets (Hangzhou Yuanda Biological Pharmaceutical), NBP capsules (Shiyao Group Enbipu Pharmaceutical), NBP reference (Shiyao Group Enbipu Pharmaceutical.), glipizide reference (Sichuan Vicchi Biochemical Technology), Soil DNA Kit (Omega Bio-Tek), Quant-iT PicoGreen dsDNA Assay Kit (ABI), First Strand cDNA Synthesis Kit (Servicebio), and SYBR Green qPCR Master Mix (Servicebio).

### 1.3 Animals

Male Sprague Dawley (SD) rats (7 weeks old, 200 ± 20 g) were purchased from Beijing Huafukang Bioscience Co. Ltd. The rats were kept in a specific pathogen-free (SPF) facility. Animal experiments were approved by the Research Ethics Committee of the Second Hospital of Hebei Medical University.

## 2 Methods

### 2.1 Quantification of NBP in plasma by HPLC-MS/MS

Chromatography separation was performed on a Symmetry C_18_ column (4.6 × 150 mm, 3.5 μm), and the column temperature was 40℃. The mobile phase consisted of water (A) and acetonitrile (B, containing 0.1% formic acid) with a flow rate of 0.8 mL·min^-1^. The elution procedure was as follows: 0-2.0 min, 75%-95% B; 2.0-6.5 min, 95% B; 6.5– .0 min, 95%– 75% B; and 7.0-8.0 min, 75% B. The injection volume was 5 μL, and the internal standard (IS) was glipizide.

#### 2.1.1 Mass spectrum conditions

Both NBP and glipizide were monitored in positive ESI mode. The scanning mode was multireaction monitoring (MRM), with the ion transitions of m/z 191.1→45.1 and 446.2→321.0, respectively. The declustering potential (DP) and collision energy (CE) of NBP were 60V and 22V, and those of the glipizide were 100V and 23V, respectively.

#### 2.1.2 Plasma sample processing method

Acetonitrile (150 μL) containing glipizide (250 ng·mL^-1^) was added to plasma (50 μL) and centrifuged at 10,900 g for 5 min. The supernatant was transferred to the autosampler vial for HPLC-MS/MS detection.

#### 2.1.3 Standard curves

A series of NBP solutions with a concentration of 20-2000 ng·mL^-1^ were added to the blank plasma to prepare the simulated plasma samples, which were tested according to the plasma sample processing method. The standard curve was drawn with the concentration of NBP as the abscissa (X) and the peak area ratio of NBP to the glipizide as the ordinate (Y). The standard curve equation was obtained through regression.

### 2.2 Experimental design and sample collection

Before the experiment, 30 SD male rats were randomly divided into three groups (n=10): the antibiotic, probiotic, and control groups. The antibiotic group received 50 mg·kg^-1^ of vancomycin solution. The probiotic group received 600 mg·kg^-1^ of live bifidobacterium tetrad bacteria suspension. The control group received physiological saline. All groups received daily gavage for 7 days. Fecal samples were collected on day 0 and day 8 before NBP administration. On day 8, all rats were gavaged using 70 mg·kg^-1^ of NBP solution. Blood samples (about 300 μL) were collected in heparinized centrifuge tubes at 0.08, 0.16, 0.33, 0.5, 0.75, 1, 1.5, 2, 3, 4, 6, and 8 h after gavage. Plasma samples were centrifuged at 8000 RPM for 10 min at 4℃. The rats were sacrificed after collecting blood and samples from the liver and small intestine tissues. The samples were stored at –80℃.

### 2.3 Effects of the intestinal microbiota on the pharmacokinetics of NBP

The frozen plasma samples were thawed naturally, processed according to the “plasma sample processing method,” and injected for analysis. NBP concentrations at different time points were calculated according to the standard curve equation. The pharmacokinetic parameters were calculated using the DAS 2.0 software. Statistical analysis was performed with SPSS 20 software to investigate the effect of antibiotic or probiotic interventions on NBP pharmacokinetics.

### 2.4 CYP3A1 mRNA and protein expressions in the liver and small intestinal

#### 2.4.1 Quantitative real-time PCR (qRT-PCR) to detect CYP3A1 mRNA expression

The frozen small intestine and liver tissue samples were naturally thawed at room temperature. Total RNA was extracted and tested for RNA concentration and purity, followed by reverse transcription and primer amplification (primer sequence shown in **Table 1**), with three replicates per transcript. CYP3A1 was encoded by *Cyp3a1*, and its internal reference was β-actin. Data processing was performed using the 2^-ΔΔCt^ method.

**Table 1.**
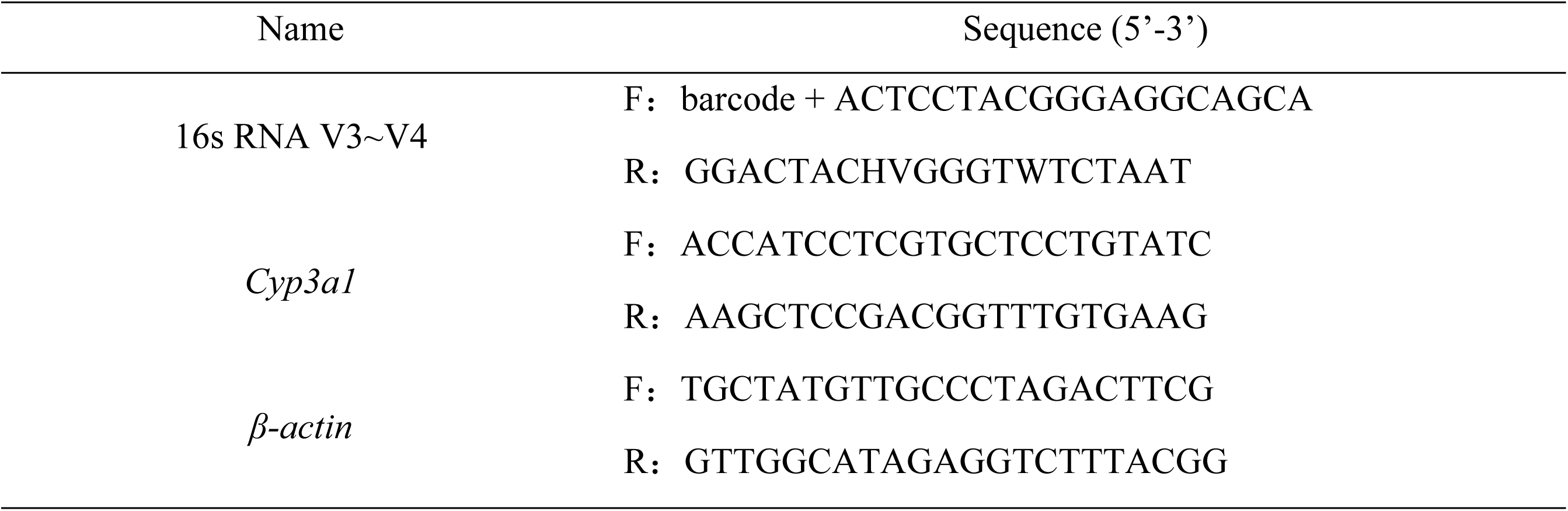
The sequence of primers.

#### 2.4.2 Western Blot assay for CYP3A1 protein expression

The frozen small intestine and liver tissue samples were thawed at room temperature. CYP3A1 total protein was extracted, added to 10% SDS-PAGE gel, and then transferred to the nitrocellulose membranes. Nitrocellulose membranes were incubated with the monoclonal CYP3A4 antibody (1: 2000 dilution) at 4℃ for 12 h, followed by the incubation with a secondary antibody (1:5000 dilution) at room temperature for 30 min. The GAPDH was used as a loading control. The Odyssey CLx Imaging System was used to detect target proteins. The protein level was normalized to the control group and expressed as multiple changes relative to the control group.

### 2.5 Effects of antibiotics and probiotics on the intestinal microbiota

The frozen feces were naturally thawed at room temperature, and DNA was extracted. PCR amplification was performed on specific primers for the V3-V4 region of 16S rRNA bacterial genes according to the Illumina 16S metagenomic sequencing library (**Table 1**). The amplified products were quantified and mixed for library construction and quality inspection. After the individual quantification step, amplicons were pooled in equal amounts and subjected to paired-end 2 × 250 bp sequencing to understand the effects of antibiotics and probiotics on the intestinal microbiota.

### 2.6 Data analysis and statistics

16S rRNA intestinal microbiota data were analyzed using the Genescloud cloud platform online. The relative abundance of the microbiota was plotted as a stacked-bar plot at the phylum and genus levels. α-diversity indices (diversity within a sample) were calculated using filtered data. Among these indices, the Chao1 diversity index estimated total richness, and the Shannon diversity index evaluated the richness and evenness of species at the genus level. β-diversity (sample dissimilarity, the difference between two samples) was calculated using the principal coordinate analysis (PCoA) based on the Bray-Curtis dissimilar matrix. Kruskal-Wallis rank sum test was used to analyze bacteria with significant abundance differences between groups, and linear discriminant analysis effect size (LEfSe) was used to analyze bacteria enrichment.

Microbial functional abundances based on marker gene sequences were predicted by Phylogenetic Investigation of Communities by Reconstruction of Unobserved States (PICRUSt2) in the Kyoto Encyclopedia of Genes and Genomes (KEGG). Spearman correlation analysis was performed to explore the relationship between the intestinal microbiota and the metabolic pathways of NBP. *P*<0.05 was considered statistically significant.

## 3 Results

### 3.1 Effects of the intestinal microbiota on the pharmacokinetics of NBP

The concentration-time curve shows that the standard curve equation for NBP in the plasma of SD rats is Y=0.00359X+0.000347 (*r*=0.9995) (**Figure 1**). **Table 2** shows the pharmacokinetic parameters of NBP. Compared to the control group, in the antibiotics group, the values of C_max_, AUC_0-8_, AUC_0-∞_, and t_1/2_ increased by 56.1% (*P<*0.001), 56.4% (*P<*0.001), 53.2& (*P*<0.001), and 24.4% (*P*<0.05), respectively. In contrast, the CL and T_max_ values decreased by 57.1% (*P<*0.001) and 28.6% (*P*<0.05), respectively. There were no statistically significant changes in pharmacokinetic parameters between the probiotic and control groups.

**Figure 1.**
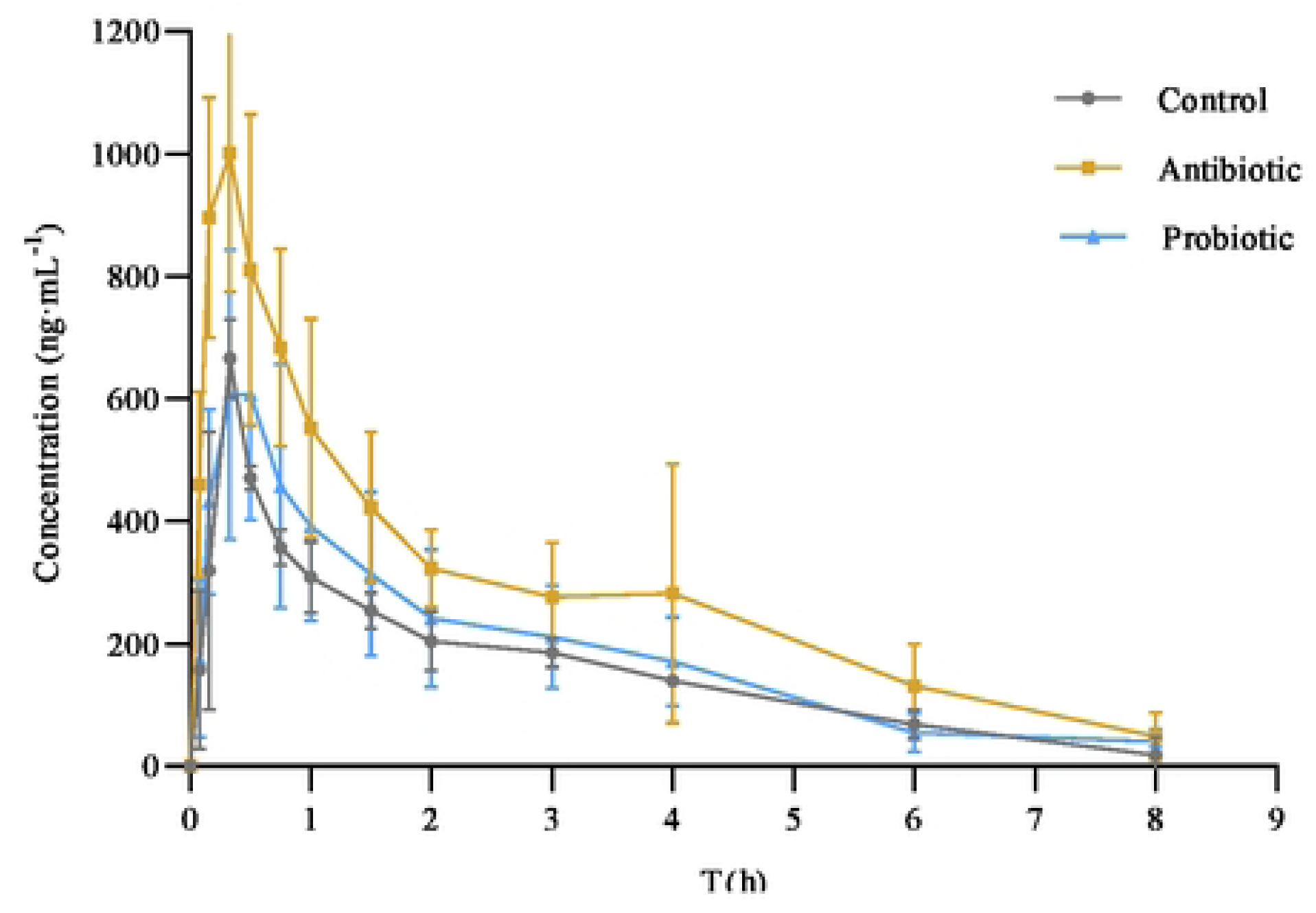
Blood concentration-time curve of n-butylphthalide in rats

**Table 2.** Pharmacokinetic parameters of n-butylphthalide in rats (mean±SD, n=10)

| Parameters | Control group | Antibiotic group | Probiotic group |
| --- | --- | --- | --- |
| $t_{1/2}(h)$ | 1.19±0.20 | 1.48±0.32* | 1.14±0.19 |
| $C_{\max}(\text{ng}\cdot\text{mL}^{-1})$ | 679.18±181.54 | 1059.30±199.58*** | 785.05±202.43 |
| $T_{\max}(h)$ | 0.35±0.05 | 0.25±0.08* | 0.35±0.25 |
| $AUC_{0-8}(\text{ng}\cdot\text{mL}^{-1}\cdot h)$ | 1327.73±254.23 | 2077.68±299.40*** | 1414.92±429.69 |
| $AUC_{0-\infty}(\text{ng}\cdot\text{mL}^{-1}\cdot h)$ | 1425.16±301.59 | 2183.88±280.66*** | 1601.98±495.69 |
| CL/$F(\text{mL}/\text{min})$ | 0.07±0.02 | 0.03±0.01*** | 0.06±0.01 |

\* $P<0.05$ , \*\* $P<0.01$ , \*\*\* $P<0.001$ , compared to the control group

### 3.2 CYP3A1 mRNA and protein expressions in the liver and small intestine

The effects of antibiotics and probiotics on CYP3A1 mRNA and protein expressions in SD rats were investigated by qRT-PCR and Western Blot (**Figure 2**). CYP3A1 mRNA and protein expression values in the small intestine of the antibiotic group were 61.2% and 66.1% of those of the control group (**Figure 2A**), respectively. The CYP3A1 mRNA and protein expression values in the liver were 44.6% and 63.9% of those of the control group (**Figure 2B**), respectively. However, there were no significant changes in CYP3A1 mRNA and protein expressions between the probiotic and control groups.

**Figure 2.**
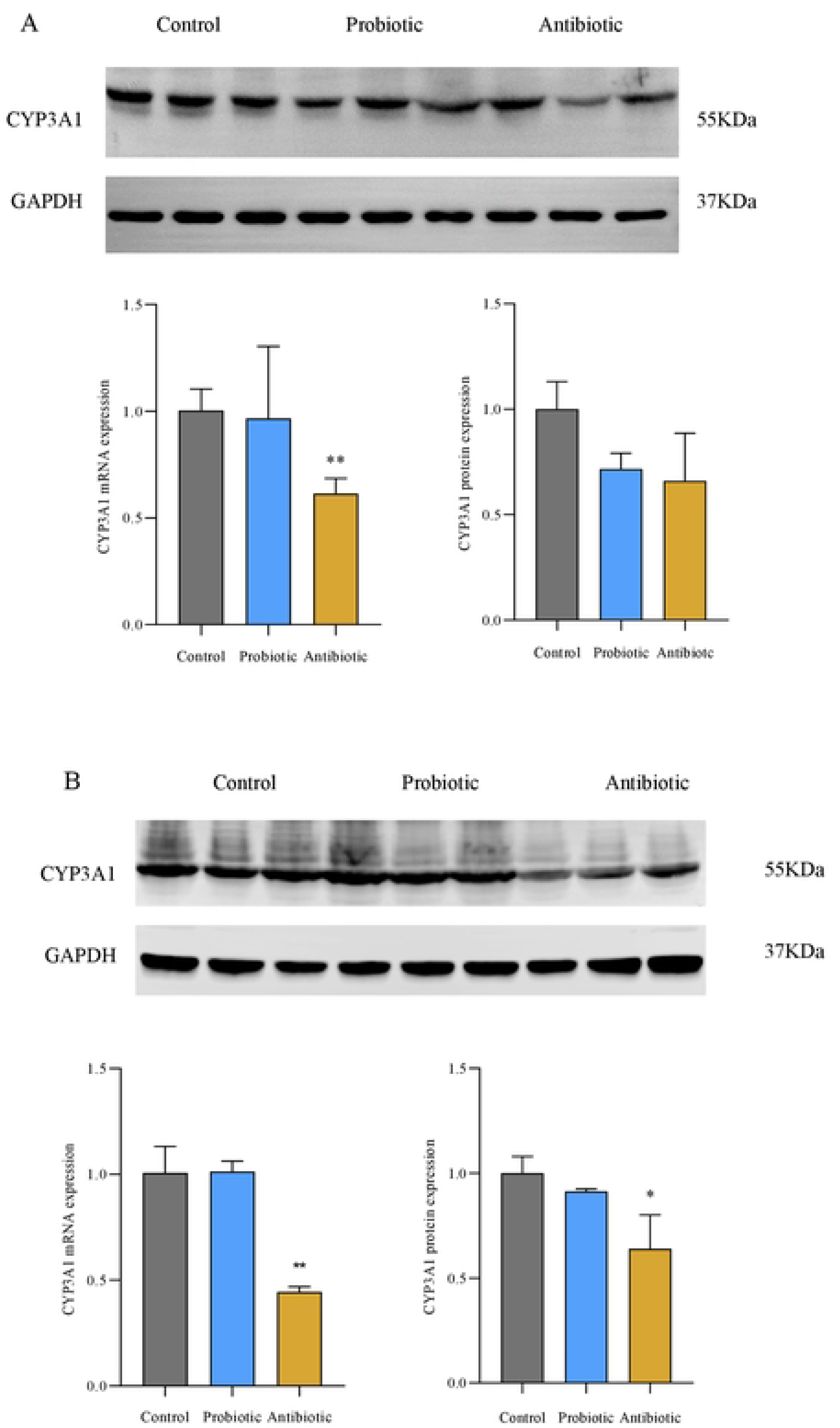
The CYP3A1mRNA and protein expressions in the small intestine and the liver (n=3) A: CYP3A1 expression in the small intestine; B: CYP3A1 expression in the liver; \**P*<0.05, \*\**P*<0.01, compared to the control group

### 3.3 Effects of antibiotics and probiotics on the intestinal microbiota

The intestinal microbiota was affected by intragastric administration of antibiotics and probiotics (**Figure 3**). There were differences in the relative abundance of the microbiota at the phylum level between the control and the antibiotic groups **(Figure 3A**). In the antibiotic group, the relative abundance of *Firmicutes* and *Bacteroidetes* decreased with increasing *Proteobacteria* and *Spirillum*. In the probiotic group, the relative abundance of *Firmicutes* increased with decreased *Bacteroidetes*. At the intestinal microbiota genus level, the dominant bacteria in the control and probiotic groups are *Lactobacillus* and *Prevotella* (**Figure 3B**). In contrast, the prevalent bacteria in the antibiotic group are *Lactobacillus* and *Sphaerochaeta.* The relative abundance of the *Lactobacillus* decreased significantly compared to the control group, and *Prevotella* was basically exhausted.

**Figure 3.**
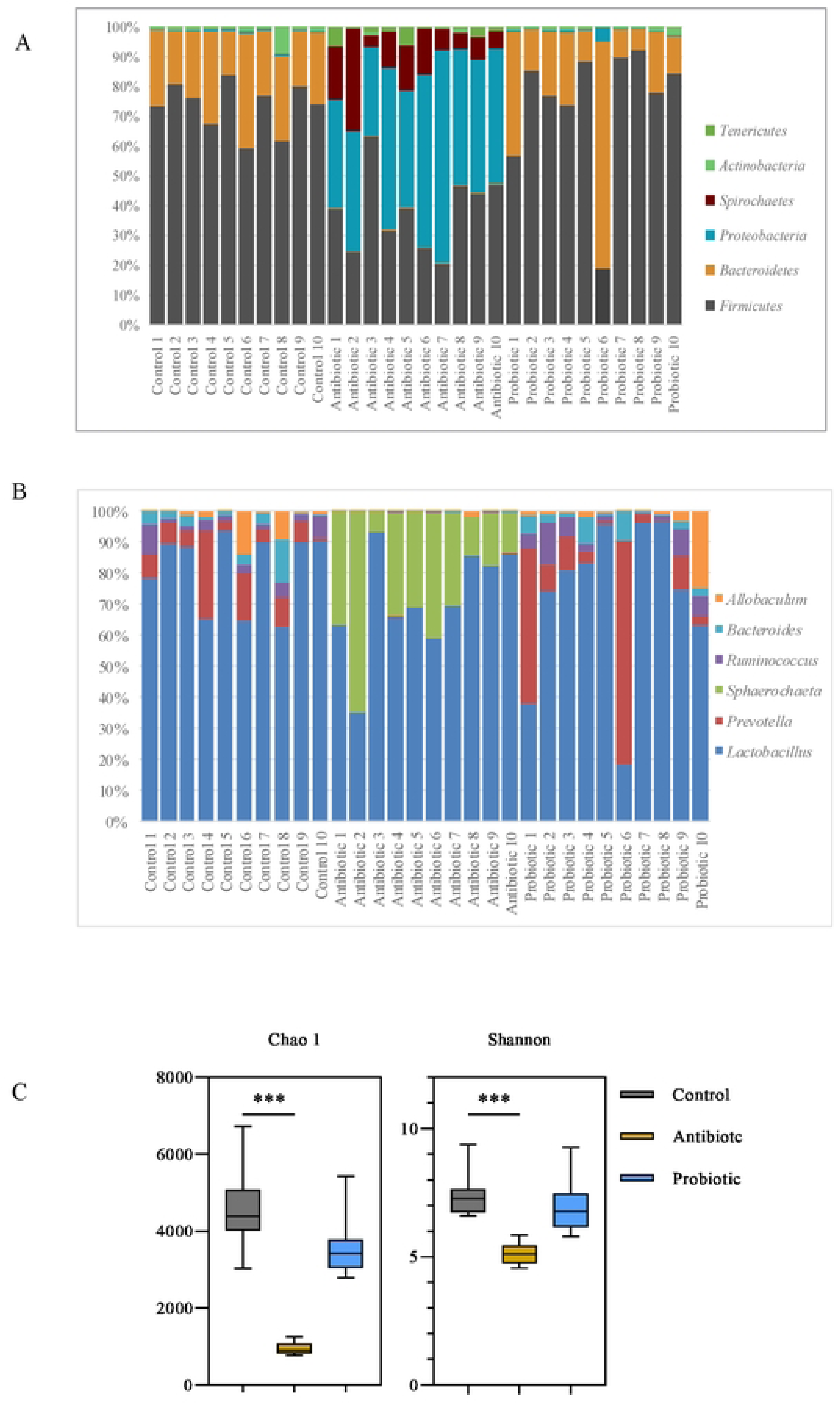
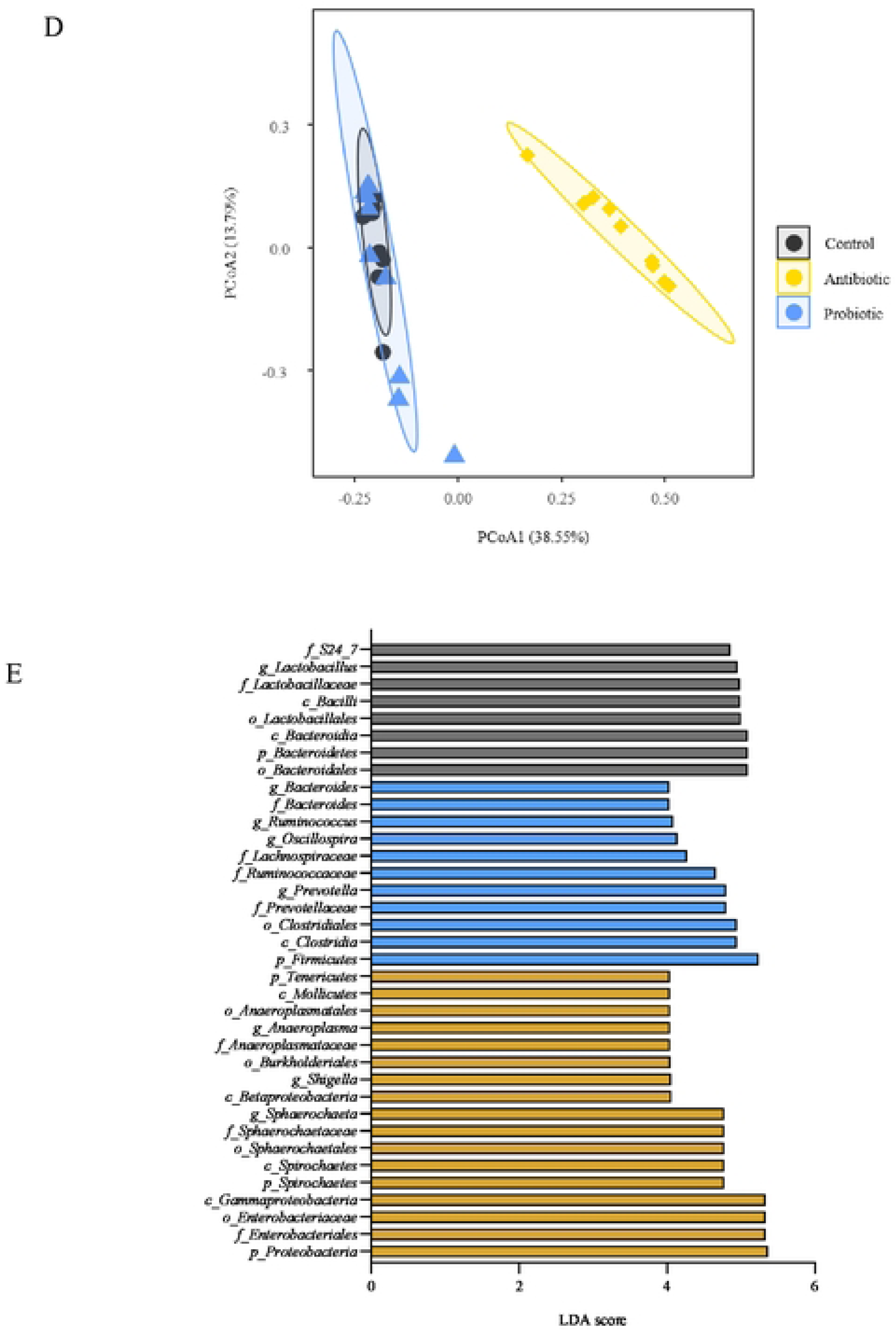
Effects of antibiotics and probiotics on intestinal microbiota. A: Phylum level of intestinal microbiota; B: Genus level of intestinal microbiota; C: α diversity; D: β diversity; E: Analysis of marker species; \*\*\**P*<0.001

Compared to the control group, the richness and evenness of bacteria in the antibiotic group decreased significantly based on α-diversity indices (*P*<0.001). In contrast, the changes in the probiotic group were not statistically significant (**Figure 3C**). The PCoA plot shows a discriminated β-diversity between the control and antibiotic groups (**Figure 3D**). LEfSe results show that *Sphaerochaeta, Shigella, Anaeroplasma,* and *Sutterella* are the signature species in the antibiotic group and were significantly enriched. In contrast, *Prevotella*, *Oscillospira*, *Allobaculum*, and *Ruminococcus* are the predominant species in the probiotic group and were significantly increased (**Figure 3E**, *P*<0.05, LDA ≥ 4.0).

### 3.4 Correlation between the intestinal microbiota and the pharmacokinetic parameters of NBP and CYP3A1 expression

The relative abundance of 20 genera showed a correlation between the pharmacokinetic parameters and the CYP3A1 expression (**Figure 4**). The relative abundance of *Anaeroplasma*, *Paenibacillus*, *and Sphaerochaeta* is positively correlated with AUC_0-8_ or C_max_ but negatively correlated with CYP3A1 expression. *Bacteroides*, *Prevotella*, *Clostridium*, and *Ruminococcus* have a positive correlation with CL and CYP3A1 mRNA expressions. Among the four genera of bacteria supplemented with probiotics, only *Bifidobacterium* is both negatively correlated with AUC_0-8_ but positively correlated with CYP3A1 mRNA expression in the small intestine. The CYP3A1 protein expression in the liver was negatively regulated by *Oscillospira*.

**Figure 4.**
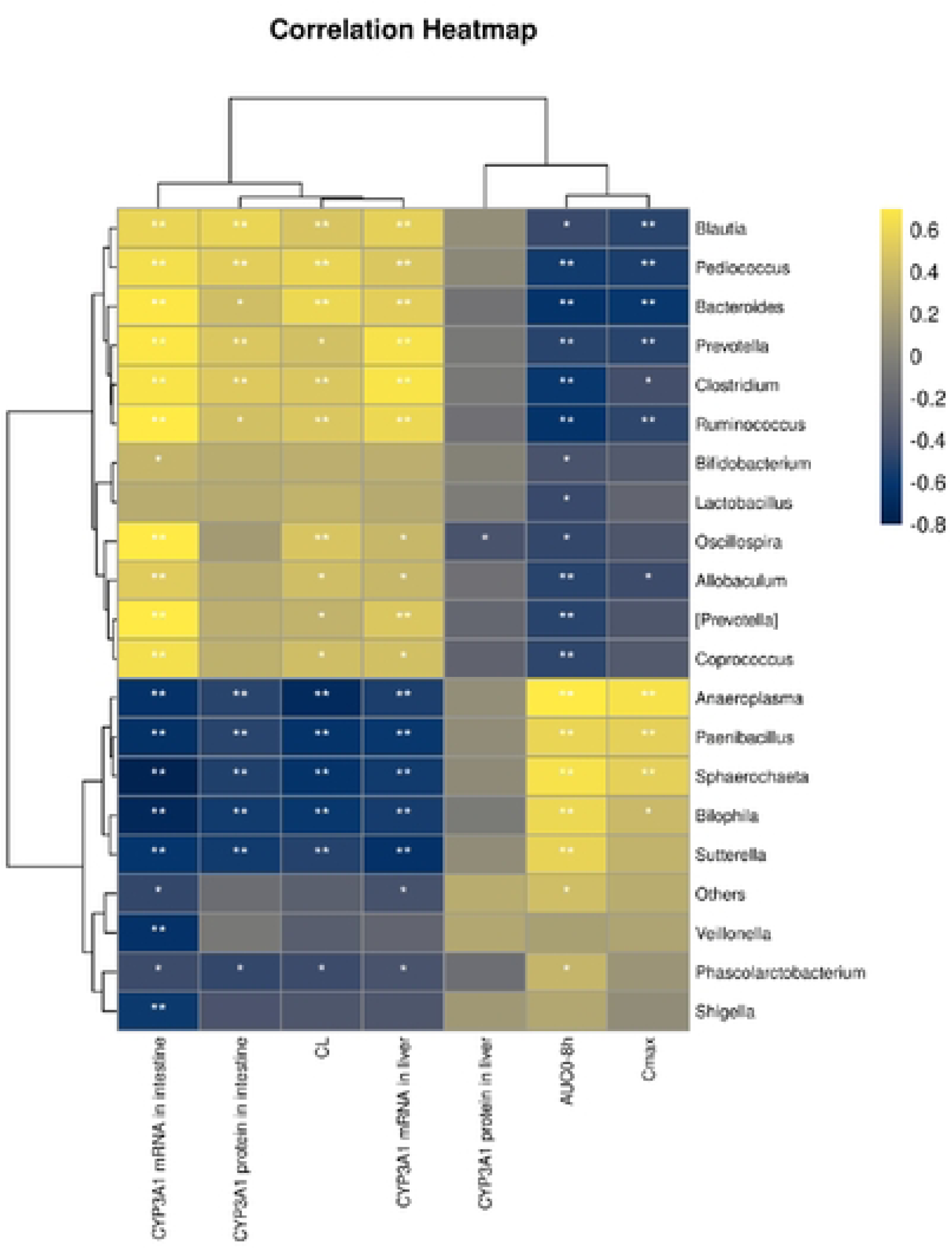
Correlation of the horizontal abundance of the intestinal microbiota genus with the pharmacokinetic parameters of n-butylphthalide and the expression of CYP3A1. \**P*<0.05; \*\**P*<0.01

### 3.5 Effects of intestinal microbiota alteration on intestinal metabolic pathways

KEGG analysis shows the effects of antibiotics and probiotics on intestinal metabolic pathways (**Figure 5**). Changes in the intestinal microbiota had the greatest influence on the KEGG primary pathway (L1) of *Metabolism* (**Figure 5A**). Analysis of the KEGG secondary pathway (L2) in the *Metabolic* pathway shows that “*Carbohydrate metabolism,”* “*Glycan biosynthesis and metabolism,”* “*Lipid metabolism,”* “*Metabolism of cofactors and vitamins,”* and “*Xenobiotics biodegradation and metabolism”* pathways were inhibited in the antibiotic group compared to the control group (**Figure 5B**, *P*<0.001). Analysis of the KEGG level 3 pathway (L3) shows no significant differences in enrichment pathways in the probiotic group. In contrast, there were 37 significant differences in the antibiotic group (Figure 5C, *P*<0.05, LDA ≥ 3).

**Figure 5.**
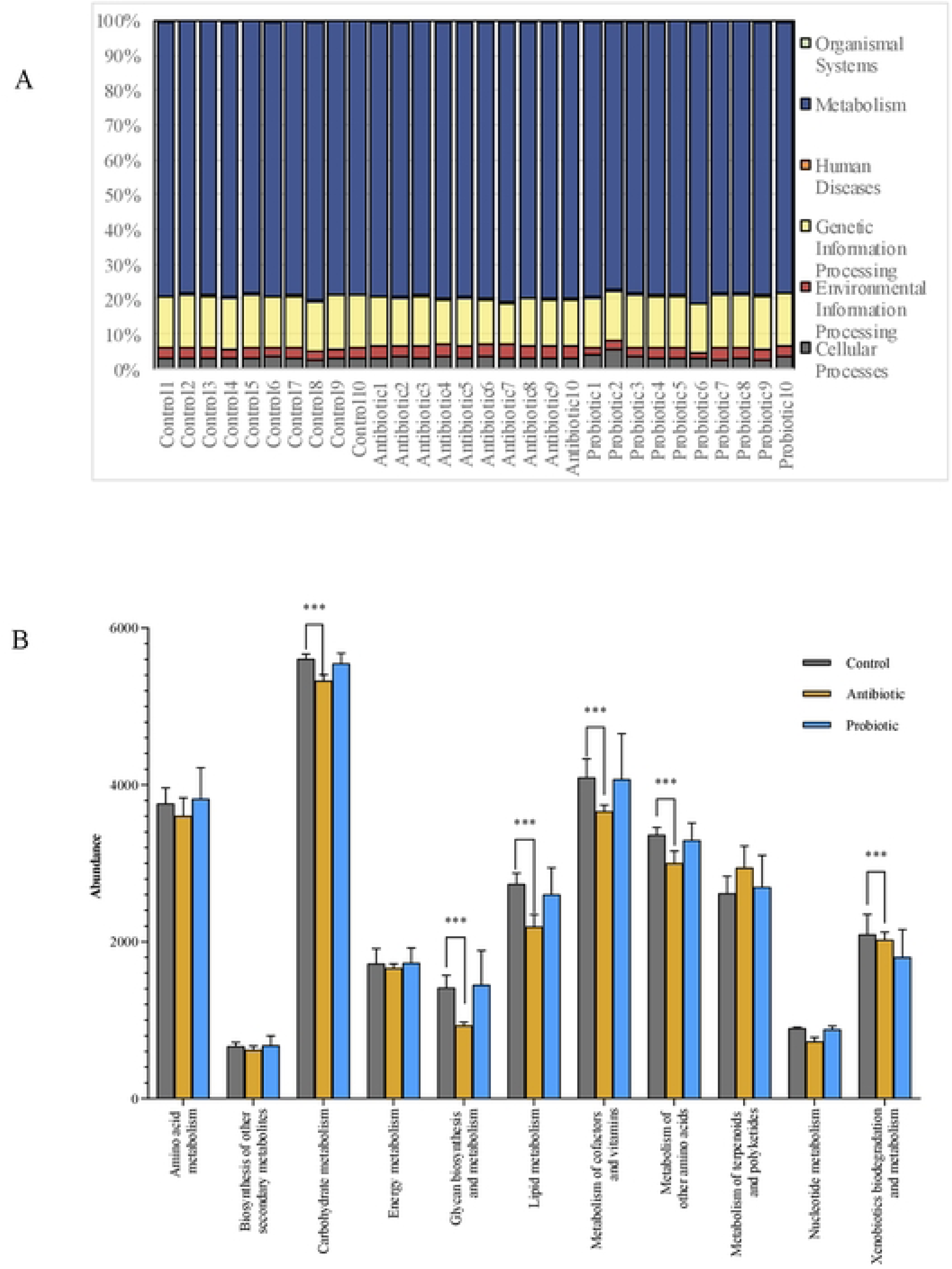
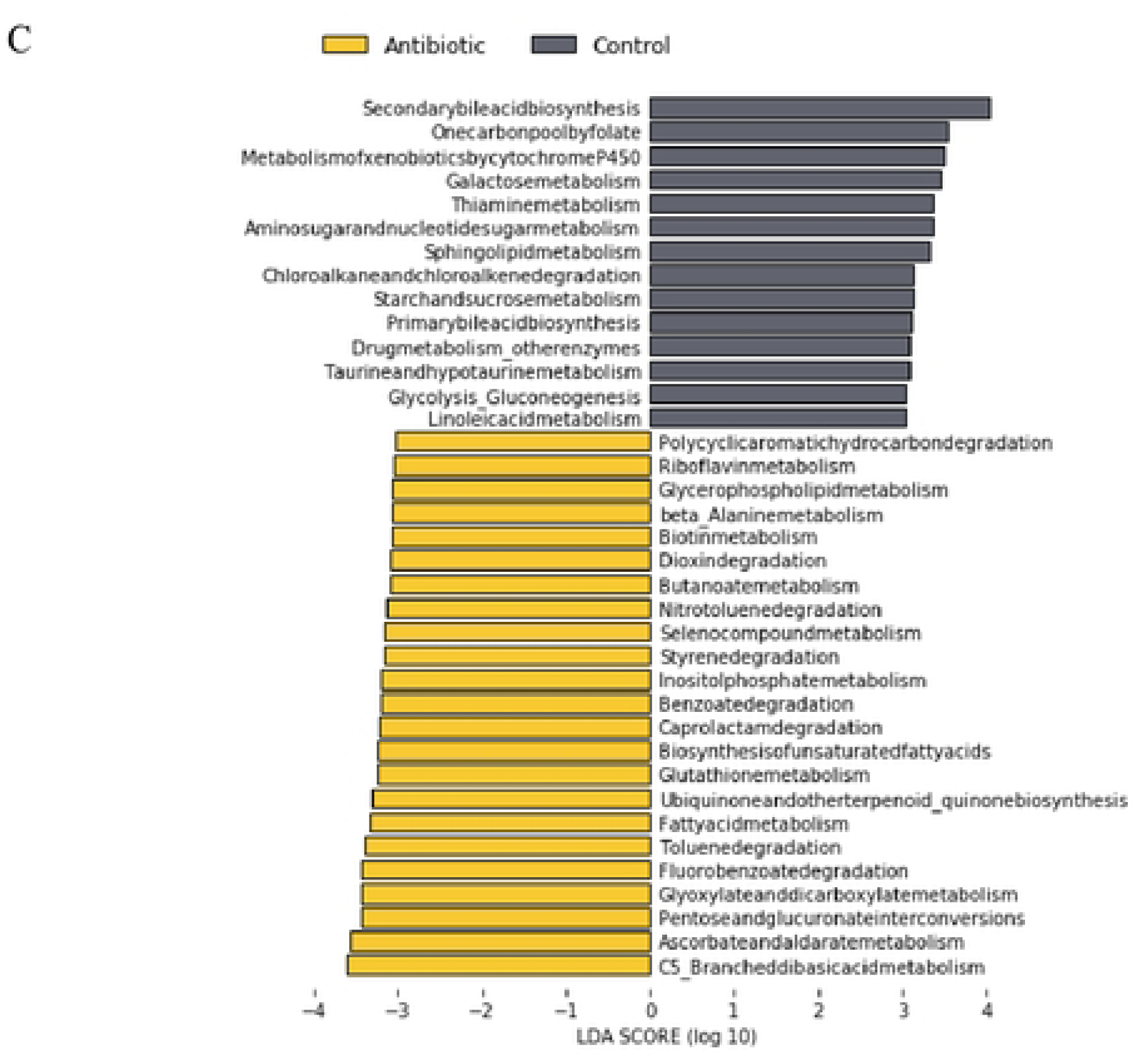
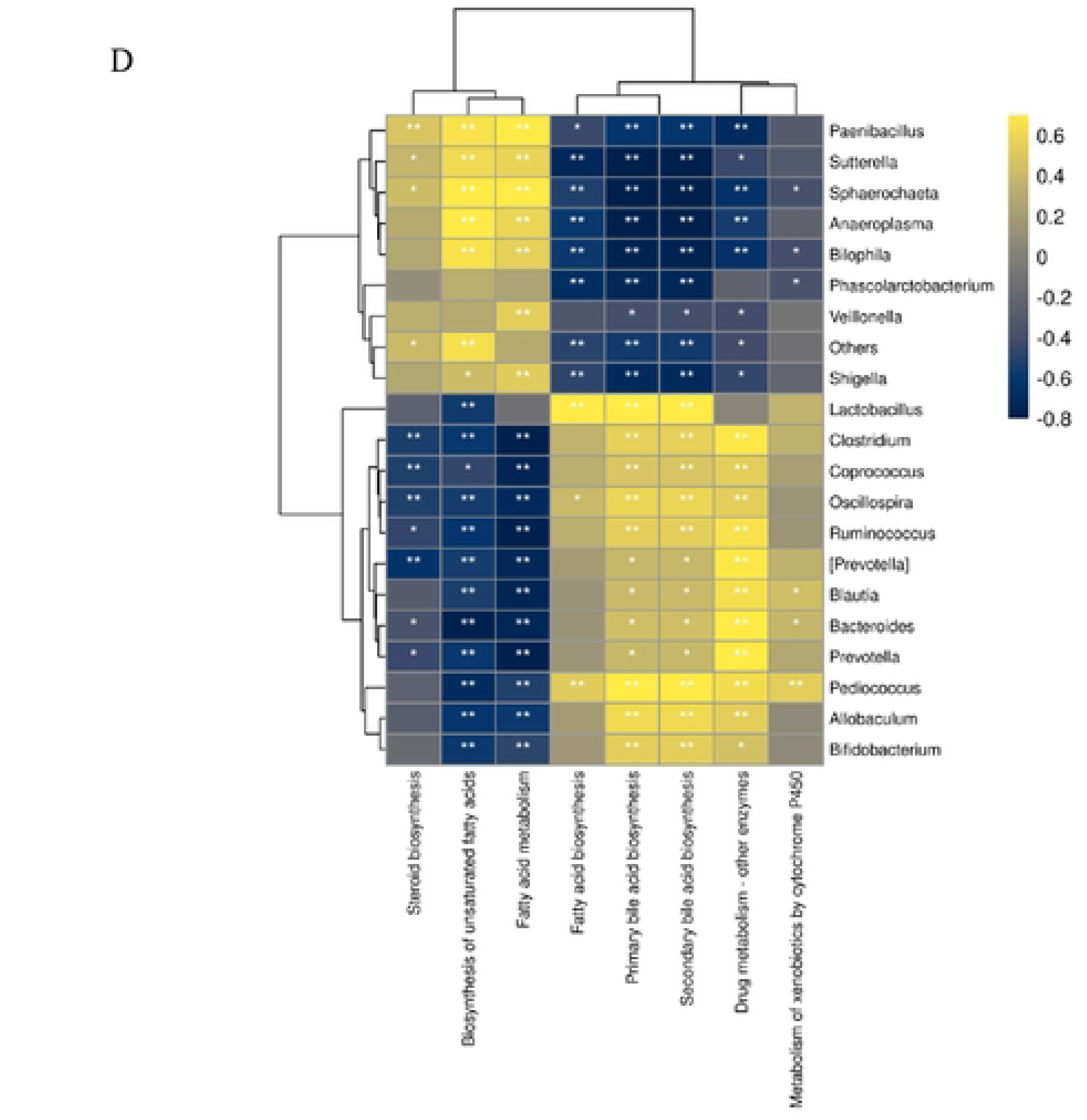
Effects of the intestinal microbiota on metabolic pathways. A: Effect of the intestinal microbiota on the KEGG L1 pathway; B: Abundance comparison of KEGG L2 metabolic pathways; C: Comparison of significantly enriched KEGG L3 pathways; D: Correlation between horizontal abundance and the KEGG L3 pathway; \**P*<0.05; \*\**P*<0.01; \*\*\**P*<0.001

Twenty relatively abundant genera were selected to analyze the correlation between the intestinal microbiota and the metabolic pathways of NBP (**Figure 5D**). The *Sphaerochaeta*, *Shigella*, *Anaeroplasma,* and *Sutterella* antibiotic group marker genera were negatively correlated with “*Secondary bile acid biosynthesis,”* “*Drug metabolism-other enzyme,”* and “*Fatty acid biosynthesis.”* In contrast, these genera were positively associated with “*Fatty acid metabolism*” and “*Biosynthesis of unsaturated fatty acids*.” These correlations were reversed for *Lactobacillus, Prevotella*, and *Oscillospira*. The “*Metabolism of xenobiotics by cytochrome P450”* was positively correlated with the relative abundance of *Pediococcus*.

## 4 Discussion

In recent years, as pharmacomicrobiomics has received increasing attention [13], the impact of intestinal flora changes on drug pharmacokinetics and pharmacodynamics has become a research hotspot [14–16]. CYP3A4 is the main metabolic enzyme of NBP in humans, and CYP3A1 in SD rats is the homologous enzyme of human CYP3A4 [17]. Therefore, this study explored the effects of intestinal flora environments on NBP pharmacokinetics and CYP3A1 expressions by administering antibiotics and probiotics to SD rats.

### 4.1 Effects of intestinal microbiota on the pharmacokinetics of drugs

Probiotics can have various beneficial effects on human health [18]. However, when combined with drugs, probiotics can change drug pharmacokinetics through several possible mechanisms. First, the intestinal flora secretes a variety of phase I and phase II metabolic enzymes, which can directly participate in the metabolism of oral drugs and affect the efficacy and toxicity of drugs [19–21]. Second, the alteration of the intestinal microbiota influences the expression of liver drug metabolism enzymes [22, 23]. Secondary bile acids and short-chain fatty acids (SCFAs), the metabolites of intestinal flora, can enter the liver through the portal vein and alter mRNA expression of multiple drug enzymes [24–27].Furthermore, the intestinal microbiota can also affect intestinal drug transporters [28, 29], maintain intestinal structure and mucosal integrity, and alter drug absorption [30].

### 4.2 Effects of antibiotic treatment on intestinal microbiota and NBP metabolism

Vancomycin was chosen as the interventional antibiotic in this study because it is not absorbed orally. It has a high intestinal concentration and a marked inhibitory effect on certain intestinal bacteria, leading to substantial changes in the intestinal microbiota [31]. In addition, vancomycin does not induce host metabolic enzymes. It does not have drug-drug interactions mediated by metabolic enzymes and is suitable for studying the pharmacokinetics of NBP as an antibiotic for intestinal flora intervention.

Lithocholic acid, one of the pregnane X receptors (PXR) ligands, is produced mainly by the normal intestinal dominant phyla *Bacteroides* and *Firmicutes*, which can up-regulate CYP3A1 mRNA expressions and increase the metabolism of CYP3A1 substrates [32–34]. Another PXR ligand, indole-3-propionic acid (IPA) [35], is the active product of tryptophan. *Clostridium* mediates the conversion of tryptophan to IPA and up-regulates the CYP3A1 mRNA expression. This study demonstrates that antibiotic gavage decreases the relative abundance of *Bacteroidetes* and *Firmicutes*, among them Clostridium, leading to a down-regulation of PXR-regulated CYP3A1 expression.

Previous studies have demonstrated that depletion of *Prevotellaceae* and enrichment with *Shigella* reduce intestinal butyrate production and downregulate the expression of host CYP450 enzymes [36]. The present study showed that in the antibiotic group, the relative abundance of *Lactobacillus* and *Prevotella* decreased, while the relative abundance of *Sphaerochaeta* and *Shigella* increased. The expression of CYP3A1 mRNA in the small intestine was significantly down-regulated, and CYP3A1 mRNA and protein expressions in the liver were significantly decreased. The results are consistent with previous research.

The results of PICRUSt2 showed that antibiotic-induced changes in the microbiota could inhibit metabolism-related pathways, thus increasing the bioavailability of NBP. The relative abundance of probiotics *Lactobacillus, Prevotella, Bacteroides,* and *Oscillospira* decreased in the antibiotic group, and metabolic pathways such as “*secondary bile acid biosynthesis*,” “*fatty acid biosynthesis*,” and “*drug metabolism*” were significantly down-regulated. The relative abundance of *Pediococcus* was positively correlated with intestinal “*metabolism of xenobiotics by CYP450*”. This suggests that intestinal microbiota imbalance, metabolic disturbance, and antibiotic-induced decreased production of metabolites such as secondary bile acids and SCFAs are the main reasons for the increased oral bioavailability of NBP.

### 4.3 Effects of probiotic gavage on intestinal microbiota and NBP metabolism

The probiotics used in this study were a mixture of four live bacteria. Among them, *Bifidobacterium*, *Lactobacillus,* and *Enterococ* are the normal flora of the human intestinal tract, which can grow, reproduce, and colonize the intestinal tract after oral administration. They can inhibit some pathogenic bacteria in the intestinal tract, maintain normal intestinal peristalsis, and regulate the balance of intestinal flora. *Bacillus* can create an anaerobic environment in addition to the normal flora in the human intestinal tract and promote the growth and reproduction of anaerobic bacteria such as *Bifidobacteria*. Previous studies have shown that the effects of bifidobacterial tetrads on the host are mainly mediated by metabolites rather than by modulation of bacterial diversity [37].

This study showed that probiotic treatment did not cause significant changes in intestinal microbiota diversity, consistent with previous studies. After supplementation with probiotics, the relative abundance of *Bifidobacterium*, *Lactobacillus,* and *Oscillospira* increased, which can promote the production of SCFAs [38]. However, there were no significant differences in the expressions of CYP3A1 and KEGG metabolic pathways compared to the control group, which may be related to individual differences [39]. In previous studies [40, 41], the effect of changes in the intestinal microbiota on drug pharmacokinetics was not consistent because differences in the type, dose, and treatment cycle of probiotics may have different effects on drug pharmacokinetics.

## 5. Conclusions

This study demonstrates that antibiotic-induced alterations in the intestinal microbiota can increase the oral bioavailability of NBP. However, CYP3A1 expression and metabolic pathways did not show significant differences in the probiotic group compared to the control group. Therefore, further studies designed to observe the effect of single probiotic supplementation on drug metabolism are warranted. Furthermore, the relationship between pharmacokinetics and pharmacodynamics is complex. Additional validation is needed to determine whether changes in NBP pharmacokinetics caused by antibiotic-mediated alterations in the intestinal microbiota affect its clinical efficacy.

## Acknowledgments

The authors thank Dong Weichong, Li Wenli and other colleagues for participation in this study.

## References

1. Singh V, Sadler R, Heindl S, et al. The gut microbiome primes a cerebroprotective immune response after stroke. J Cereb Blood Flow Metab. 2018 Aug; 38(8): 1293–1298.

2. Benakis C, Poon C, Lane D, et al. Distinct commensal bacterial signature in the gut is associated with acute and long-term protection from ischemic stroke. Stroke. 2020 Jun; 51(6): 1844–1854.

3. 18. Benakis C, Brea D, Caballero S, et al. Commensal microbiota affects ischemic stroke outcome by regulating intestinal γδ T cells. Nat Med. 2016 May; 22(5): 516–523.

4. Mulroy E, Bhatia KP. The gut microbiome: A therapeutically targetable site of peripheral levodopa metabolism. Mov Disord Clin Pract. 2019 Aug 26; 6(7): 547–548.

5. Walsh J, Gheorghe CE, Lyte JM, et al. Gut microbiome-mediated modulation of hepatic cytochrome P450 and P-glycoprotein: impact of butyrate and fructo-oligosaccharide-inulin. J Pharm Pharmacol. 2020 Aug; 72(8): 1072–1081.

6. Banoth S, Tangutur AD, Anthappagudem A, et al. Cloning and *in vivo* metabolizing activity study of CYP3A4 on amiodarone drug residues: A possible probiotic and therapeutic option. Biomed Pharmacother. 2020 Jul; 127: 110128.

7. Zhou J, Ouyang J, Gao Z, et al. MagMD: Database summarizing the metabolic action of gut microbiota to drugs. Comput Struct Biotechnol J. 2022 Nov 12; 20: 6427–6430.

8. Cussotto S, Walsh J, Golubeva AV, et al. The gut microbiome influences the bioavailability of olanzapine in rats. EBioMedicine. 2021 Apr; 66: 103307.

9. Diao X, Deng P, Xie C, et al. Metabolism and pharmacokinetics of 3-n-butylphthalide (NBP) in humans: the role of cytochrome P450s and alcohol dehydrogenase in biotransformation. Drug Metab Dispos. 2013 Feb; 41(2): 430–44.

10. Jia J, Wei C, Liang, et al. The effects of DL-3-n-butylphthalide in patients with vascular cognitive impairment without dementia caused by subcortical ischemic small vessel disease: A multicentre, randomized, double-blind, placebo-controlled trial. Alzheimers Dement. 2016 Feb; 12(2): 89–99.

11. Fan X, Shen W, Wang L, et al. Efficacy and safety of DL-3-n-Butylphthalide in the treatment of poststroke cognitive impairment: a systematic review and meta-analysis. Front Pharmacol. 2022 Jan 25; 12: 810297.

12. Houlden A, Goldrick M, Brough D, et al. Brain injury induces specific changes in the caecal microbiota of mice via altered autonomic activity and mucoprotein production. Brain Behav Immun. 2016 Oct; 57: 10–20.

13. Doestzada M, Vila AV, Zhernakova A, et al. Pharmacomicrobiomics: a novel route towards personalized medicine? Protein Cell. 2018 May; 9(5): 432–445.

14. Weersma RK, Zhernakova A, Fu J. Interaction between drugs and the gut microbiome. Gut. 2020 Aug; 69(8): 1510–1519.

15. Wu B, Chen M, Gao Y, et al. *In vivo* pharmacodynamic and pharmacokinetic effects of metformin mediated by the gut microbiota in rats. Life Sci. 2019; 226: 185–192.

16. Jarmusch AK, Vrbanac A, Momper JD, et al. Enhanced characterization of drug metabolism and the influence of the intestinal microbiome: a pharmacokinetic, microbiome, and untargeted metabolomics study. Clin Transl Sci. 2020; 13: 972–984.

17. Martignoni M, Groothuis GM, de Kanter R. Species differences between mouse, rat, dog, monkey and human CYP-mediated drug metabolism, inhibition and induction. Expert Opin Drug Metab Toxicol. 2006 Dec; 2(6): 875–94.

18. O’Toole PW, Marchesi JR, Hill C. Next-generation probiotics: the spectrum from probiotics to live biotherapeutics. Nat Microbiol. 2017 Apr 25; 2: 17057.

19. Kiang TK, Ensom MH, Chang TK. UDP-glucuronosyltransferases and clinical drug-drug interactions. Pharmacol Ther. 2005 Apr; 106(1): 97–132.

20. Zhang X, Han Y, Huang W, et al. The influence of the gut microbiota on the bioavailability of oral drugs. Acta Pharm Sin B. 2021 Jul; 11(7): 1789–1812.

21. Drozdzik M, Busch D, Lapczuk J, et al. Protein abundance of clinically relevant drug-metabolizing enzymes in the human liver and intestine: a comparative analysis in paired tissue specimens. Clin Pharmacol Ther. 2018 Sep; 104(3): 515–524.

22. Walsh J, Gheorghe CE, Lyte JM, et al. Gut microbiome-mediated modulation of hepatic cytochrome P450 and P-glycoprotein: impact of butyrate and fructo-oligosaccharide-inulin. J Pharm Pharmacol. 2020 Aug; 72(8): 1072–1081.

23. Wang S, Wen Q, Qin Y, et al. Gut microbiota and host cytochrome P450 characteristics in the pseudo germ-free model: co-contributors to a diverse metabolic landscape. Gut Pathog. 2023 Mar 21; 15(1): 15.

24. Avior Y, Levy G, Zimerman M, et al. Microbial-derived lithocholic acid and vitamin K2 drive the metabolic maturation of pluripotent stem cells-derived and fetal hepatocytes. Hepatology. 2015 Jul; 62(1): 265–278.

25. Mun SJ, Lee J, Chung KS, et al. Effect of Microbial short-chain fatty acids on CYP3A4-mediated metabolic activation of human pluripotent stem cell-derived liver organoids. Cells. 2021 Jan 11; 10(1): 126.

26. Selwyn FP, Cheng SL, Klaassen CD, et al. Regulation of Hepatic Drug-Metabolizing Enzymes in Germ-Free Mice by Conventionalization and Probiotics. Drug Metab Dispos. 2016 Feb; 44(2): 262–74.

27. Banoth S, Tangutur AD, Anthappagudem A, et al. Cloning and *in vivo* metabolizing activity study of CYP3A4 on amiodarone drug residues: A possible probiotic and therapeutic option. Biomed Pharmacother. 2020 Jul; 127: 110128.

28. Kaddurah-Daouk R, Baillie RA, Zhu H, et al. Enteric microbiome metabolites correlate with response to simvastatin treatment. PLoS One. 2011; 6(10): e25482

29. Li H, Wang J, Fu Y, et al. The bioavailability of glycyrrhizinic acid was enhanced by probiotic *Lactobacillus rhamnosus* R0011 supplementation in liver fibrosis rats. Nutrients. 2022 Dec 11; 14(24): 5278

30. Natividad JM, Agus A, Planchais J, et al. Impaired aryl hydrocarbon receptor ligand production by the gut microbiota Is a key factor in metabolic syndrome. Cell Metab. 2018 Nov 6; 28(5): 737–749.e4.

31. Haak BW, Lankelma JM, Hugenholtz F, et al. Long-term impact of oral vancomycin, ciprofloxacin and metronidazole on the gut microbiota in healthy humans. J Antimicrob Chemother. 2019 Mar 1; 74(3): 782–786.

32. Bilen M, Dufour JC, Lagier JC, et al. The contribution of culturomics to the repertoire of isolated human bacterial and archaeal species. Microbiome. 2018 May 24; 6(1): 94.

33. Singh V, Lee G, Son H, et al. Butyrate producers, “the sentinel of gut”: their intestinal significance with and beyond butyrate, and prospective use as microbial therapeutics. Front Microbiol. 2023 Jan 12; 13: 1103836.

34. Boesmans L, Valles-Colomer M, Wang J, et al. Butyrate producers as potential next-generation probiotics: safety assessment of the administration of *butyricicoccus pullicaecorum* to healthy volunteers. mSystems. 2018 Nov 6; 3(6): e00094–18.

35. Wikoff WR, Anfora AT, Liu J, Schultz PG, et al. Metabolomics analysis reveals large effects of gut microflora on mammalian blood metabolites. Proc Natl Acad Sci U S A. 2009 Mar 10; 106(10): 3698–703.

36. Chen Y, Liu Y, Wang Y, et al. Prevotellaceae produces butyrate to alleviate PD-1/PD-L1 inhibitor-related cardiotoxicity via PPARα-CYP4X1 axis in colonic macrophages. J Exp Clin Cancer Res. 2022 Jan 3; 41(1): 1.

37. Bai T, Xu Z, Xia P, et al. The Short-term efficacy of bifidobacterium quadruple viable tablet in patients with diarrhea-predominant irritable bowel syndrome: potentially mediated by metabolism rather than diversity regulation. Am J Gastroenterol. 2022 Dec 23.

38. Yang J, Li Y, Wen Z, et al. Oscillospira – a candidate for the next-generation probiotics. Gut Microbes. 2021 Jan-Dec; 13(1): 1987783.

39. Cussotto S, Walsh J, Golubeva AV, et al. The gut microbiome influences the bioavailability of olanzapine in rats. EBioMedicine. 2021 Apr; 66: 103307.

40. Liu J, Cheng Y, Zhang Y, et al. Lactobacillus rhamnosus induces CYP3A and changes the pharmacokinetics of verapamil in rats. Toxicol Lett. 2021 Nov 1; 352: 46–53.

41. Lee HJ, Zhang H, Orlovich DA, et al. The influence of probiotic treatment on sulfasalazine metabolism in rat. Xenobiotica. 2012 Aug;42(8):791–7.

